# CFTR High Expresser BEST4+ cells are pH-sensing neuropod cells: new implications for intestinal physiology and Cystic Fibrosis disease

**DOI:** 10.1101/2025.01.24.634747

**Authors:** Diego C. dos Reis, Jason Jin, Anderson Santos, Caroline Muiler, Eleanor Zagoren, Martin Donnelley, David Parsons, Patricia Cmielewski, Nicole Reyne, Alexandra McCarron, Zachary Smith, Kaelyn Sumigray, Nadia A. Ameen

## Abstract

Single-cell RNA sequencing (scRNA-seq) studies identified a novel subpopulation of epithelial cells along the rostrocaudal axis of human intestine specifically marked by bestrophin 4 (BEST4) that are enriched for genes regulating pH, GPCR acid-sensing receptors, satiety, cGMP signaling, HCO3^-^ secretion, ion transport, neuropeptides, and paracrine hormones. Interestingly, BEST4+ cells in the proximal small intestine express CFTR but have not been linked to the previously described CFTR High Expresser Cell (CHE) subpopulation in rat and human intestine. ScRNA-seq studies in rat jejunum identified CHEs and a gene expression profile consistent with human small intestinal BEST4+ and neuropod cells. Protein immunolocalization confirmed that CHEs express CFTR, BEST4, neuropod proteins, high levels of intracellular uroguanylin (UGN), guanylyl cyclase-C (GC-C), and the proton channel otopetrin 2 (OTOP2), and display long basal processes connecting to neurons. OTOP2, GC-C, and CFTR traffic robustly into the apical domain of CHEs in response to acidic luminal conditions, indicating their roles in luminal pH regulation. In the ΔF508 cystic fibrosis (CF) rat jejunum, the loss of apical CFTR did not affect BEST4 protein expression in CHEs. However, there was an increased abundance of CHE cells in the ΔF508 rat jejunum compared to wild-type animals. Furthermore, ΔF508 rat CHEs expressed higher levels of GC-C at the apical domain compared to wild-type. These data implicate CHEs in intestinal CF disease pathogenesis.

**NEW & NOTEWORTHY:** This is the first study to identify CFTR High Expresser cells in the rat small intestine as neuropod cells capable of sensing and responding to luminal pH. This study also provides the first characterization of CFTR and relevant mRNA and proteins in CHEs in CF rat models that provide insights into the significance of CHEs to CF intestinal disease.

## INTRODUCTION

CFTR High Expresser cells (CHEs) constitute a rare subpopulation of enterocytes, found in the rat and human intestine, initially identified through in situ hybridization (ISH) studies of the cystic fibrosis transmembrane conductance regulator (*CFTR*) mRNA (1, 2). Subsequently, our group confirmed very high levels of CFTR at the protein level using immunolabeling techniques and designated these cells as CHEs (3, 4). CHEs are sporadically localized in the rat and human duodenum and jejunum but distinctly absent in mice (3, 5), represent 1-3% of total enterocytes, and are found predominantly on the villi but also in the superficial crypts excluding the stem cell compartment (6). Notably, they display prominent mitochondria within their cytoplasm, prompting us to compare them to mitochondria-rich (MR) cells and ionocytes of various species, including amphibians, fish, sea turtles, and higher mammals (4, 7).

Previous protein characterization studies unveiled a unique expression profile of ion channels and transporters at the apical and basolateral membranes of villus CHEs. Villus CHEs are enriched for the basolateral Na^+^K^+^2Cl^-^-cotransporter 1 (NKCC1) that regulates intestinal secretion, vacuolar-ATPase proton pump (V-ATPase), NHE regulatory factor (NHERF1), Na^+^/K^+^-ATPase, and syntaxin 3 but lack absorptive hydrolases and other ion transporters found on the brush border of mature villus enterocytes (3, 6). Although ultrastructural studies confirm that CHEs are enterocytes on the villi, the brush border is immature, disordered and shorter than neighboring enterocytes (4). Furthermore, stimulation with second messengers cAMP, cGMP, calcium, or acetylcholine known to activate CFTR-mediated secretion, leads to robust trafficking of CFTR from subapical vesicles into the brush border of villus CHEs, implying a pivotal role in fluid secretion in an absorptive epithelium (14,15).

Single-cell RNA sequencing (scRNA-seq) studies have introduced a new avenue for mapping and elucidating the relevance of newly identified rare populations of cells in the normal and diseased intestine. In this context, several groups identified a novel population of intestinal cells expressing bestrophin 4 (*BEST4*) throughout the small and large intestine in many species (excluding mice) that are enriched for genes known to be important in satiety, fluid and HCO_3_^-^ transport and pH regulation including carbonic anhydrase 7 (*CA7*), guanylate cyclase activator 2A (*GUCA2A*), guanylate cyclase activator 2B (*GUCA2B*), and otopetrin 2 (*OTOP2)* (*8*). Interestingly, segment-specific expression of gene transcripts in BEST4+ cells point to functional specialization of this cell population along the rostrocaudal axis of the intestine. Consistent with this, BEST4+ cells in the small intestine fall into the secretory lineage, while those in the colon are mature absorptive colonocytes (9). Indeed, BEST4+ cells in the proximal small intestine (but not colon) express CFTR implicating a specific role in secretion and cystic fibrosis (CF) intestinal disease pathophysiology (8–10). Additionally, transcriptomic analyses in human intestine revealed that BEST4+ cells are enriched for the neuropeptides vasoactive intestinal peptide (*VIP*) and neuropeptide Y (*NPY*), suggesting a potential role in neuropod functionality (9).

Based on the high expression of *BEST4* in these cells throughout the small and large intestine, they are referred to as BEST4+ cells. BEST4 is a member of the bestrophin family of calcium (Ca^2+^)-dependent chloride (Cl^-^) ion channels highly permeable to HCO_3_^-^ at the basolateral membrane of epithelial cells. Bestrophins have been studied in retinal epithelial cells, but little is known about the role of BEST4 in the intestine (11, 12). Interestingly, like our observations of CHEs, BEST4 is not observed in the crypt stem cell compartment and its expression increases as cells undergo maturation (3, 6, 9). *BEST4* mRNA was first identified in human duodenal CHEs, which were then referred to as BCHE cells (13). Although detailed ultrastructural protein localization and trafficking studies on rat CHEs have been published (3, 4, 6, 14, 15), scRNA-seq studies have largely failed to connect BEST4+ cells in the small intestine to CHEs.

The current study utilized data from scRNA-seq analysis of epithelial cells in the rat jejunum to identify CHE-specific enriched gene transcripts. Antibodies were used to label CHE-specific proteins identified in the scRNA-seq analysis to provide further insights into the physiological and pathophysiological relevance of CHEs. We confirmed that rat CHEs were positive for BEST4 and enriched with acid sensing receptors, proton channels, proteins regulating cGMP signaling, CFTR-mediated HCO_3_-secretion, and neuropod genes. Immunolocalization studies identified long basal processes in CHEs, filled with vesicular BEST4 and UGN, the functional protein of the *Guca2b* gene, connecting to neural structures in the lamina propria (LP). Immunolabeling for neuropod cell markers confirmed that CHEs express neuropod cell proteins. Since little is known about CHEs in the CF intestine, we examined changes in CFTR and CHE-specific proteins implicated in pH regulation and HCO_3_^-^ secretion under conditions of luminal acidity and in the jejunum of a CF rat model expressing the most prevalent human CF disease mutation Phe508del (ΔF508). Our data confirm that rat CHEs express high levels of BEST4 and support a role for CHEs as novel neuropod cells that sense and respond to luminal acidity in the proximal small intestine by rapid communication with the enteric nervous system (ENS).

## MATERIALS AND METHODS

### scRNA-seq

One Gene Expression Omnibus Series (GSE) (GSE272055) was obtained from NCBI (National Center for Biotechnology Information) and GEO (Gene Expression Omnibus). Differentially expressed genes (DEGs) were processed for GSEA (Gene Set Enrichment Analysis). CHE-related genes were identified from the existing literature, following which the relationship between Differentially Expressed Genes (DEGs) and CHE-related genes was examined.

### Antibodies

Anti-rodent CFTR antibody (#AME4991) was produced by Dr. Ameen and has been characterized previously (6, 16, 17). Anti-BEST4 antibody (#R12038) was produced and provided by Dr. Sumigray. Antibodies against MEIS1 (#MA5-27191) and OTOP2 (#PA5-62727) were purchased from Thermo Fisher. Antibodies against UGN (#6910, #6912) were provided by Dr. Michael Goy. TUBB3 (#NB100-1612), TUBB2B (#NBP2-46250), SYT3 (#NBP1-19320), and S100A6 (#NB110-93274SS) were purchased from Novus Biologicals. Additional antibodies include: MYO1B (#HPA013607, Atlas Antibodies), GC-C (#ab225864, Abcam), CHGA (#ab254557, Abcam), NFM (#CH22106, Neuromics), and ChAT (#AB143, Chemicon). Secondary antibodies were acquired from Jackson ImmunoResearch (#711-545-152, Alexa Fluor 488 Affini-Pure^TM^ Donkey Anti-Rabbit IgG; #703-295-155, Rhodamine Red^TM^-X Affini-Pure^TM^ Donkey Anti-Chicken IgY (IgG); and #715-605-151, Alexa Fluor 647 Affini-Pure^TM^ Donkey Anti-Mouse IgG). F-actin labeling was performed with Texas Red^TM^-X Phalloidin (#T7471, Invitrogen). Nuclear counterstaining was achieved via 4’,6-diamidino-2-phenylindole (DAPI) (#D1306, Invitrogen).

### Animals

The Institutional Animal Care and Use Committee of Yale University School of Medicine approved the study. Normal male Sprague-Dawley rats (200-250g, Charles River Laboratories, Wilmington, MA) were fasted overnight but allowed free access to drinking water. Rats were anesthetized with Inactin (thiobutabarbital) hydrate (120 mg/kg) (#T133, Sigma Aldrich) administered by intraperitoneal (IP) injection. Body temperature was maintained during the studies using a heating pad.

### CF Rat Tissues

CF rat breeding and tissue collection was performed under approval of the University of Adelaide Animal Ethics Committee (M-2017-010). Rat proximal jejunum tissues from a CF model carrying the ΔF508 *CFTR* mutation, *CFTR* knockout (CFKO) model, and control wild type (18, 19) were kindly provided by Dr. Martin Donnelley.

### Treatment of Rat Ligated Small Intestinal Loops and Luminal pH Measurement

Normal rat intestinal loops (2.5 cm length) were created with ligatures in the duodenum and proximal jejunum. Intestinal loops were infused with 0.2 ml sterile saline (NaCl 0.9%, pH 7.4) or saline acidified with HCl (pH 2.0) for 30 minutes (min). The abdomen was closed, and the animals were kept warm. Immediately following treatment, luminal samples (100-200 µl) were collected, and pH, Cl^-^, and HCO_3_^-^ were measured using the i-STAT blood gas analyzer (Abaxis, Union City, CA) equipped with EC8+ sample cartridges (Abbot Point of Care, Princeton, NJ) as described previously (15). Each pH, Cl^-^, and HCO_3_^-^ data point was collected in at least 3 animals. At the end of the experiment, the animals were euthanized by administration of Inactin (200 mg/kg) in accordance with Yale IACUC approval.

### Treatment of Rat Ligated Small Intestinal Loops with heat-stable enterotoxin (STa) from *Escherichia coli*

The jejunum was identified, intestinal loops (5 cm length) created with ligatures, and each loop instilled with either 0.5 ml of freshly prepared STa (0.5 M) or saline. The abdomen was closed, the animal was kept warm, and the jejunal loops were examined for fluid accumulation 30 min later as described (20). At the end of the experiment, the animals were euthanized by administration of Inactin (200 mg/kg) in accordance with Yale IACUC approval. Tissues were removed and prepared for immunolocalization studies.

### Rat Intestinal Organoid Culture

Normal rat intestinal organoids were generated and maintained according to Zagoren et al. (21). The culture medium consisted of Advanced DMEM/F-12 (#12634010, Gibco) supplemented with 20% fetal bovine serum (FBS; #16-000-044, Gibco) and 2 mM L-glutamine (#A2916801, Gibco), along with the following components: 10 nM gastrin (#G9145, Sigma), 1 mM N-acetylcysteine (#A9165-5G, Sigma), 10% Noggin-conditioned medium (#6057-NG-100, R&D Systems), 50 ng/mL recombinant mouse epidermal growth factor (EGF; #PMG8041, Thermo Fisher), 100 ng/mL recombinant human insulin-like growth factor 1 (IGF-1; #B356441, BioLegend), 100 ng/mL recombinant human fibroblast growth factor 2 (FGF2; #100-18B-250ug, Peprotech), 10% R-spondin 1 (Rspo1)-conditioned medium (#3710-001-01, R&D Systems), and 145% L-Wnt3a-conditioned medium (#CRL-2647, ATCC).

### pHrodo^TM^ Red AM Assay

Normal rat intestinal organoids were passaged and allowed to grow for 2-3 days in Matrigel. They were released from Matrigel by pipetting and incubated in 10 µM pHrodo^TM^ Red AM in 10 mM HEPES (pH7.4) for 30 minutes. Organoids were plated in 10 mM HEPES (pH 7.4) or calibration buffers (pH 4.5, 5.5, 6.5) from an Intracellular pH Calibration Buffer Kit (#P35379, Thermo Fisher) in an 8 chambered coverslip (Ibidi, 80806) for 10 minutes at 37 °C with 5% CO_2_. Images were acquired using a HC Fluotar L Visir 25X/0.95 NA water immersion objective on a Leica STELLARIS 5 Confocal Microscope with a white light laser and HyD S detectors (excitation at 560 nm, emission at 590 nm). A standard curve was generated by plotting fluorescence intensity against pH from calibration buffers, allowing for the determination of intracellular pH in HEPES relative to these standards.

### MYO1B Knockdown in Rat Intestinal Organoids

For stable MYO1B knockdown (KD), pLKO.1 lentiviral short hairpin RNA (shRNA) constructs were obtained from the Sigma-Aldrich MISSION shRNA consortium (Clone ID: TRCN0000016249), and nontargeting (scramble) control shRNA (SHC002). Stable rat intestinal organoid lines were generated by lentiviral transduction by 3×10^5^ lentiviral particles. The organoids were selected with 4 µg/mL puromycin 48 hours (h) after transduction, for 3 weeks.

### Western Blotting

MYO1B KD and scrambled shRNA organoids were lysed in 1X Cell Lysis Buffer (#9803, Cell Signaling), and protein concentrations were determined using the Bradford Assay. 20 µg of protein were mixed with 1X Laemmli Buffer (#1610737EDU, Bio-Rad) containing 5% β-mercaptoethanol and heated at 95 °C for 5 min for protein denaturation. Samples were loaded onto a 7.5% Mini-PROTEAN® TGX™ Stain-Free Precast Gel (#4568024, Bio-Rad). Electrophoresis was carried out in 1X Tris-Glycine-SDS Buffer (#1610732, Bio-Rad) at 90 V for 5 min, followed by 120 V for 45 min, or until the dye front reached the bottom of the gel. Following electrophoresis, the gel was activated using the stain-free method in the Bio-Rad ChemiDoc XRS+ Imaging System. Proteins were transferred onto a Trans-Blot Turbo Mini 0.2 µm PVDF membrane (#1704150, Bio-Rad) using the Bio-Rad Turbo Transfer system (#1704150, Bio-Rad) for 10 min. The stain-free method was then applied to the PVDF membrane to determine total protein content. After transfer, the membrane was blocked in 5% non-fat dry milk in 1X TBST (blocking solution) for 1 h at room temperature (RT). The PVDF membrane was incubated with 1:500 anti-MYO1B antibody diluted in the blocking solution, overnight at 4 °C. After primary antibody incubation, the membrane was rinsed twice for 5min with 1X TBST followed by incubation with 1:4000 HRP-conjugated goat anti-rabbit IgG (#554021, BD Biosciences) in the blocking solution for 1h at RT. The membrane was then washed 5 times with TBST for 10 min each. After the final rinse, the membrane was treated with clarity western ECL substrate (#1705061, Bio-Rad) in the dark for 5 min before acquisition. Protein bands were visualized using the chemiluminescence method in a Bio-Rad ChemiDoc XRS+ Imaging System, and the exposure time was adjusted for optimal results. The band intensities were normalized by total protein content.

### Tissue Preparation

Rat intestinal segments were immediately excised and flushed with ice-cold phosphate buffered saline (PBS, pH 7.4). Tissues were cut into 2 mm-thick rings, embedded in tissue-freezing medium optimal cutting temperature (OCT) (#4583, Sakura Finetek) and cryosectioned. Some tissues were fixed in 10% neutral buffered formalin (NBF, pH 7.4) for 24 h and embedded in paraffin wax. Tissues were also fixed in 2-4% paraformaldehyde (PFA) in PBS for 30 min at RT. PFA-fixed tissues were rinsed with PBS before undergoing cryoprotection by a 5-20% sucrose (in PBS) gradient in 30 min intervals at RT. Rings were incubated in 20% sucrose overnight at 4 °C. Tissue samples were arranged in a 4-ring array and embedded in OCT. OCT-embedded blocks were immediately frozen in isopentane pre-chilled with liquid nitrogen. Blocks were stored in - 80 °C freezer until use. 5-8 μm-thick sections were cut on a Leica Cryostat, applied to Superfrost^TM^ Plus slides (#1255015, Fisher Scientific), and stored in −20 °C. Microtome sections were prepared by flushing rat jejunum with ice-cold PBS and fixing in 4% PFA in PBS overnight at 4 °C. The tissue was then embedded in 2.5% low melting agarose (#A9414, Sigma) and cut into 175 µm-thick sections using a Precisionary Compresstome© VF-310-0Z (speed: 3.5, oscillation: 6). The rings were then stored in PBS at 4 °C.

### Histopathological Analysis

Normal (wild type), ΔF508, and CFKO rat proximal jejunum tissues were fixed in 10% Neutral Buffered Formalin (NBF) for 24 h and embedded in paraffin wax. 5 μm-thick sections were prepared and deparaffinized using a xylene substitute (#A5597, Sigma) followed by rehydration through a graded ethanol series. These sections were subsequently stained with hematoxylin and eosin (H&E). Sections were examined under light microscopy to identify inflammatory cellular changes and morphological alterations consistent with CF pathology in humans.

### Immunofluorescence (IF) Labeling

OCT-embedded unfixed sections were thawed at RT for 5 min and fixed with 2-4% PFA for 3 min. Sections were washed with PBS and incubated for 1 h in blocking solution prepared with 5% normal goat serum, 5% normal donkey serum, and 0.2% Triton X-100 (#T8787, Sigma) in PBS (PBST, pH 7.4). Primary antibodies were combined into a single cocktail and applied to slides for double or triple-labeling staining. Slides were left overnight at 4 °C, rinsed with PBS, and incubated with secondary antibodies diluted in blocking solution for 1 h at RT. F-actin filaments were visualized by staining sections with Texas Red^TM^-X for 30 min at RT. Sections were incubated with DAPI for 10 min at RT before coverslip mounting with Prolong^TM^ Diamond Antifade (#P36961, Invitrogen). Slides were dried overnight at 4 °C and imaged.

Fixed tissues (2-4% PFA), were cryoprotected, embedded in OCT, and sectioned. Tissue sections were thawed at RT before hydrating with PBS. Sections were treated with 1% sodium borohydride in PBS for 20 min to minimize background autofluorescence, and then washed in PBS. IF labeling steps continued as described above for unfixed tissue. Slides were mounted with antifade, dried overnight at 4 °C, and imaged. Paraffin embedded sections were deparaffinized in xylene substitute and rehydrated through a graded ethanol series. Slides were rinsed in deionized water for 5 minutes and submitted to heat-induced epitope retrieval (HIER) using ethylenediaminetetraacetic acid (EDTA) buffer solution (pH 9.0) at 95 °C for 20 min in a water bath. After HIER, slides were cooled at RT for 20 min and washed with PBS. Primary antibodies were added into a single cocktail and applied to slides for double-labeling staining. Slides were incubated overnight at 4 °C, rinsed with PBS, and incubated with Alexa Fluor 488-conjugated anti-rabbit, and 647-conjugated anti-mouse secondary antibodies for 1 h at RT. Nuclei was stained using DAPI for 10 min at RT before coverslip mounting with antifade before drying and imaging. All antibodies were diluted in blocking solution.

Microtome sections were incubated in blocking solution at RT on a rocker for 2 h. Following blocking, primary antibody in blocking solution was added and incubated at 37 °C for 2 days. After incubation, the sections were washed 3 times, 10 min each with PBST. Secondary antibodies were then added, and the sections were incubated at 37 °C for 1 day on a rocker. The sections underwent further washing after secondary incubation, with PBST for 10 min repeated 3 times. They were then transferred to slides and Ce3D mounting medium (N-methylacetamide, #M26305, Sigma; Iohexol, #AN1002424, Accurate Chemical; and Triton X-100 in PBS) for overnight incubation at 4 °C.

The organoids were fixed with 4% PFA for 10 min at RT and washed with PBS, releasing them from Matrigel. Next, organoids were resuspended in blocking solution and incubated at RT for 45 minutes in a rocker. After the blocking step, diluted anti-CFTR antibody was added in the blocking solution and incubated on the rocker at room temperature for 45 minutes, followed by three washes in 1X PBST. Then, organoids were washed twice with PBST for 5 minutes. A secondary cocktail containing secondary antibody, phalloidin-AF647, and DAPI were added to the organoids and incubated at RT for 30 minutes. The organoids were washed 5 times for 5 min each wash and mounted with antifade Prolong^TM^ Diamond Antifade.

### Fluorescence In Situ Hybridization

Rat intestinal segments were excised and flushed with ice-cold PBS. The tissue was cut into 2mm-thick rings and fixed in 10% NBF for 24 h at RT. Rings were stored in 70% ethanol at 4 °C after fixation. Tissue samples were processed and embedded in paraffin wax, cut into 5 μm-thick sections on Superfrost^TM^ Plus slides, and stored dry at RT. The FFPE slides were deparaffinized by two consecutive incubations in xylene substitute for 5 min and dehydrated with two 95% ethanol washes for 2 min. Fluorescence in situ hybridization (FISH) was performed on sections with the RNAscope^TM^ Multiplex Fluorescence Reagent Kit v2 with TSA Vivid Dyes (#323270, ACD, Inc.) and RNA probes for *CFTR* (#531041-C2) and *GUCA2B* (#313511). The full protocol used was developed by ACD, Inc. and is available on their website (#UM323100). Sections were either single-labeled for *Cftr* or *Guca2b* or double-labeled. Sections were incubated with DAPI for 10 min after completion of FISH to label cell nuclei.

### Fluorescence Microscopy

Immunolabeled sections were imaged on the following microscopes: Zeiss Axio Imager.M2 with ApoTome.2 and captured on a Axiocam 506 (14 bits/channel, 2752×2208 pixels); Zeiss Axio Observer.Z1/7 with ApoTome.2 and captured on a Hamamatsu C11440 ORCA-Flash4.0 digital camera (16 bits/channel, 2048×2048 pixels). Digital images were taken at 20×, 40×, and 63× magnifications and at the same exposure times when comparing control and altered conditions unless noted otherwise. Images were processed using Zeiss Efficient Navigation (ZEN) 3.4 software and ImageJ (FIJI). Immunolabeled microtome sections were imaged on a Leica STELLARIS 5 Confocal Microscope with Power HyD S detectors (1024×1024×93 pixels). Images were taken at 25× magnification and processed using ImarisViewer 10.1. Scale bars are provided for each image.

### Image Quantification

#### CHE Abundance

The percentage of CHE cells versus total number of epithelial cells was analyzed in rat proximal jejunum immunolabeled sections using the ZEN 3.4 software. Five low-magnification (20×) images taken from randomly selected sections were analyzed from each animal. Apical CFTR and nuclear MEIS1 double-labeling were used to identify CHE cells. The total number of enterocytes were determined by counting nuclei labeled with DAPI. All measured values are presented as means ± SE.

#### CFTR and GC-C Apical/Subapical Ratio

Fluorescence intensity (FI) levels were measured over the apical and subapical membrane pole of CHEs in sections from five independent immunofluorescent labeled images from each animal using the ZEN 3.4 software. For selection of regions of interests, ribbon-shaped areas of 2.9 μm^2^ were drawn on apical and subapical compartments of CHEs using a circular tool. For background correction, pixel intensity values from the luminal areas next to the CHEs were subtracted. All measured values are presented for each individual CHE cell.

## RESULTS

### Characterization of CHE-specific proteins in normal rat jejunum

CHE cells express disproportionate levels of CFTR in the subapical vesicular compartment and apical membrane relative to villus enterocytes. Similar to findings in human intestine (9, 13), scRNA-seq in rat jejunum identified CHE-specific genes that were distinctly upregulated, including transcription factors, actin motors, G protein coupled receptor (GPCR) pH-sensing receptors, cGMP receptors, and ligands and cellular machinery responsible for acid-stimulated HCO_3_^-^ secretion (Fig 1*A*). A subpopulation of highly expressing CFTR epithelial cells also exist in the lung (22) referred to as pulmonary ionocytes (23, 24). Interestingly, *FOXI1* was identified as a major transcription factor regulating CFTR-rich ionocytes in the airway epithelium, but *Foxi1* was not identified in our scRNA analysis in the rat intestine. Instead, we identified the transcription factor Meis homeobox 1 (*Meis1*) to be enriched in and specific to CHEs (Fig 1*A* and *B*). Furthermore, immunolabeling of mouse trachea with antibodies against CFTR and FOXI1 confirmed the localization of pulmonary ionocytes expressing CFTR/FOXI1 but failed to localize FOXI1 in CHEs in rat jejunum (Fig 1*K* and *L*). To confirm that *Meis1* was specific to CHEs in the intestinal epithelium, we used an anti-MEIS1 antibody and found high levels of MEIS1 concentrated in the nuclei of CHE cells on the villi (Fig 1*C*) and superficial crypt (Fig 1*D*). CHE cells are most abundant on the villi (Fig 1*C*). They are also present in the transit-amplifying region of the crypt-villus axis, where cells begin to undergo differentiation (Fig 1*D*), as previously reported by our laboratory (3, 6).

**Fig 1.**
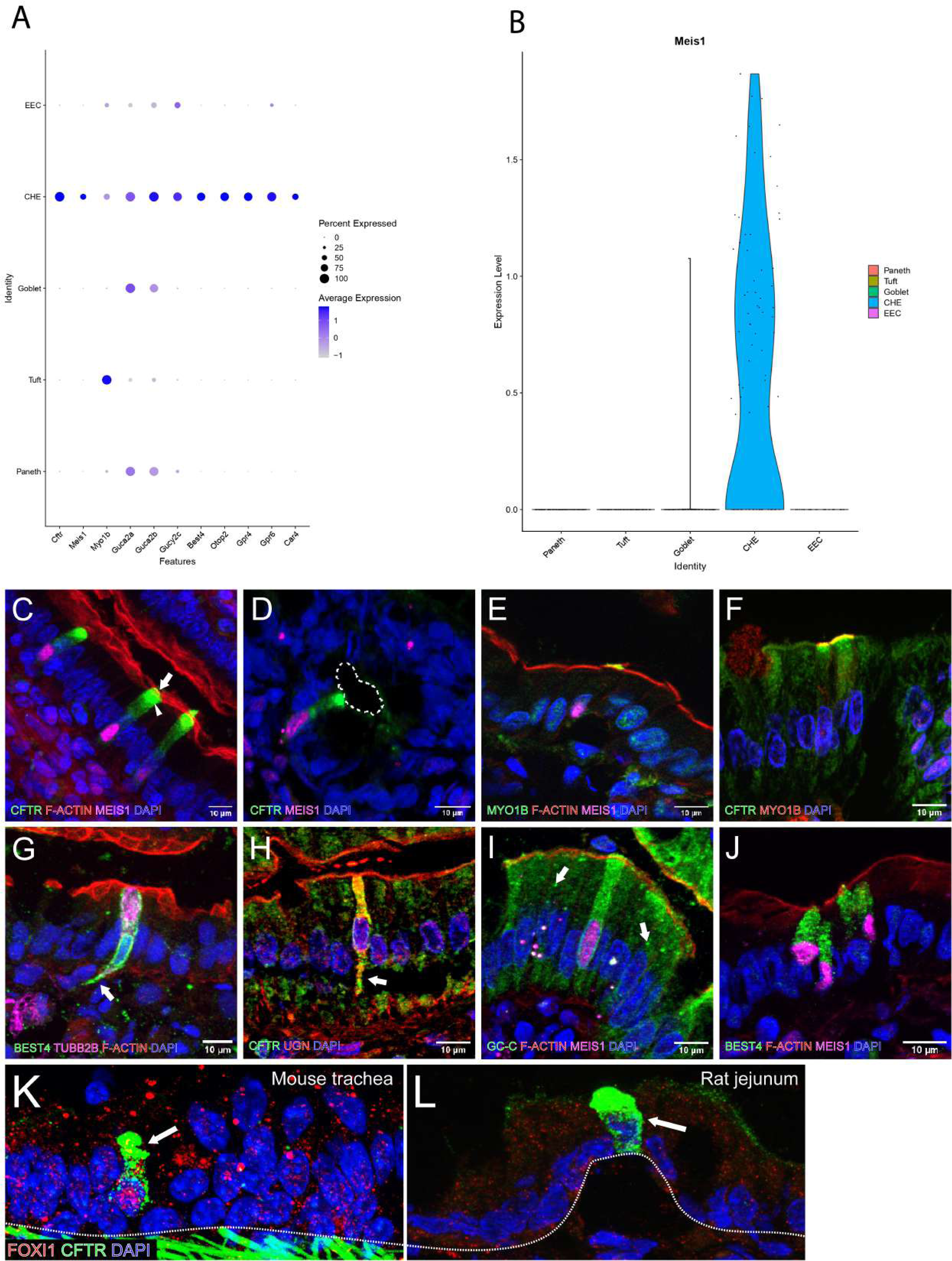
*(A)* Dot plot of CHE-specific enriched genes relative to other cell types. *(B)* MEIS1 is selectively expressed in CHEs compared to other cell types. *(C-L)* Immunolabel of CHEs in tissue sections. All regions show immunolabeled villus epithelium unless noted otherwise. MEIS1 (magenta) is a robust nuclei marker for CHE cells. *(C)* CHE cells highly express CFTR (green) in the subapical vesicular compartment *(arrowhead)* and apical membrane *(arrow)*. *(D)* CHE cells are also found in the superficial crypt region *(outline: crypt lumen). (E)* A CHE cell highly expresses MYO1B (yellow) on the apical membrane (F-actin, red). *(F)* MYO1B (red) and CFTR (green) co-localize on the apical membrane of CHE cells. *(G)* CHE cells highly express BEST4 (green) on the basolateral membrane and extends into the lamina propria in basal process *(arrow) (H)* CHE cells highly expresses cytoplasmic and apical UGN (red) and CFTR (green). All other villus epithelial cells express apical UGN (red). *(I)* A CHE cell highly expresses cytoplasmic and apical GC-C (green). *(J)* CHE cells marked by nuclear MEIS1 (magenta) are enriched with BEST4 (green) in the basal and cytoplasmic domain. *(K)* Pulmonary ionocytes in mouse trachea express high CFTR protein (green, arrow) in the apical domain and Foxi1 (red) in the nucleus *(L)* Foxi1 (red) is absent in the nucleus of CHE cells (arrow) and epithelial cells in rat jejunum. Scale bars: 10 µm.

The actin motor myosin 1A (MYO1A) is necessary for brush border development in mature villus enterocytes (25). Previous studies from our laboratory identified MYO1A as a vital motor regulating CFTR trafficking from subapical endosomes into the brush border of mature enterocytes (26). However, CHE cells display an underdeveloped brush border (4) and interestingly lack MYO1A, despite their ability to robustly traffic CFTR from subapical endosomes to the brush border (6). The motor responsible for brush border trafficking in CHEs has not been identified. scRNA-seq data identified myosin 1B (*Myo1b*) enrichment in CHEs, prompting the use of an anti-MYO1B antibody to immunolabel rat jejunum sections and determine MYO1B localization to the brush border of these cells. Indeed, MYO1B co-localized with apical F-actin (Fig 1*E*) and CFTR (Fig 1*F*), supporting its function as a crucial motor in CHE brush border trafficking. Consistent with its role in cellular traffic, ultrastructural studies of rat jejunum confirmed prominent motor fibers traversing CHE cells (4) that was further supported by abundant immunolabel for beta tubulin 2 (TUBB2) in the subapical cytoplasm (Fig 1*G*).

Human duodenal BEST4+ CHE cells express high levels of *GUCA2B* (*13*). *GUCA2B* encodes for the peptide hormone UGN, an endogenous ligand of guanylate cyclase-C (GC-C), that activates CFTR to secrete HCO_3_^-^ via cGMP signaling (27). Previous studies using the same UGN antibodies used in the current study identified immunoreactive UGN+ cells in rat jejunum that also expressed Chromogranin A (CHGA) (28). These findings led to the prevailing conclusions that the source of UGN is enteroendocrine cells (EECs) in the rat intestine (29). Interestingly, the morphological description and distribution of these UGN/CHGA+ cells in the rat jejunum are consistent with CHEs. Utilizing the same UGN antibody demonstrated by Nakazato *et al.*, we discovered that CHEs are highly enriched for UGN in the cytoplasm (Fig 1*H*) and followed the same distribution as shown previously and in Fig 3*E*. Some UGN+ cells (Fig 3*F*) and CHEs (not shown) also express CHGA. These findings are noteworthy, as our published ultrastructural studies on CHE cells in rat jejunum confirmed that they are enterocytes (4). Other studies using in situ hybridization in rat and human duodenum, and human colon, identified *Guca2b*/*GUCA2B* in solitary epithelial cells that did not express *CHGA*, suggesting that EECs are not the only source of UGN (30).

ScRNA-seq data also identified significant enrichment of *Gucy2c* (GC-C) in CHEs in the rat jejunum (Fig 1*I*). GC-C is the transmembrane receptor for UGN, guanylin (GN), and Heat Stable Enterotoxin (STa) that mediates cGMP-signaling of CFTR in the intestine (20). GC-C is expressed in epithelial cells in the small and large intestine (31, 32). Notably, recent studies in the mouse small intestine revealed GC-C enrichment in the cytoplasm of villus neuropod cells, indicating that its localization is not confined to the apical membrane (33). Indeed, immunolabeling with an antibody against GC-C in rat small intestine revealed punctate structures consistent with vesicles in the apical domain of villus enterocytes (Fig 1*I*). Additionally, GC-C was detected in punctate structures throughout the cytoplasm of CHEs, labeled with MEIS1. This distribution indicates for the first time that GC-C is present in vesicles and thus has the potential to traffic both in CHE cells and villus enterocytes.

ScRNA-seq data confirmed that human duodenal CHE cells express *BEST4*, a potential Cl^-^, HCO_3_^-^, or voltage-gated Ca^2+^ channel (13). Interestingly, CFTR is expressed solely in BEST4+ cells in the proximal small intestine but not BEST4+ colonocytes (8, 9). We identified *Best4* enrichment in rat CHEs in our scRNA seq studies, and using MEIS1 as a marker for CHEs, we confirmed that BEST4 protein is highly enriched and specific to rat CHE cells (Fig 1*J*). BEST4 is a basolateral membrane channel, but our localization suggests that, like CFTR and other ion channels, BEST4 is also present in vesicles and located in the basal domain and cytoplasm of CHE cells. Immunoreactive BEST4 is also sometimes observed in long basal pseudopodia that extend into the lamina propria (LP) (Fig 1*G* and Fig 2*A* and *B*).

**Fig 2.**
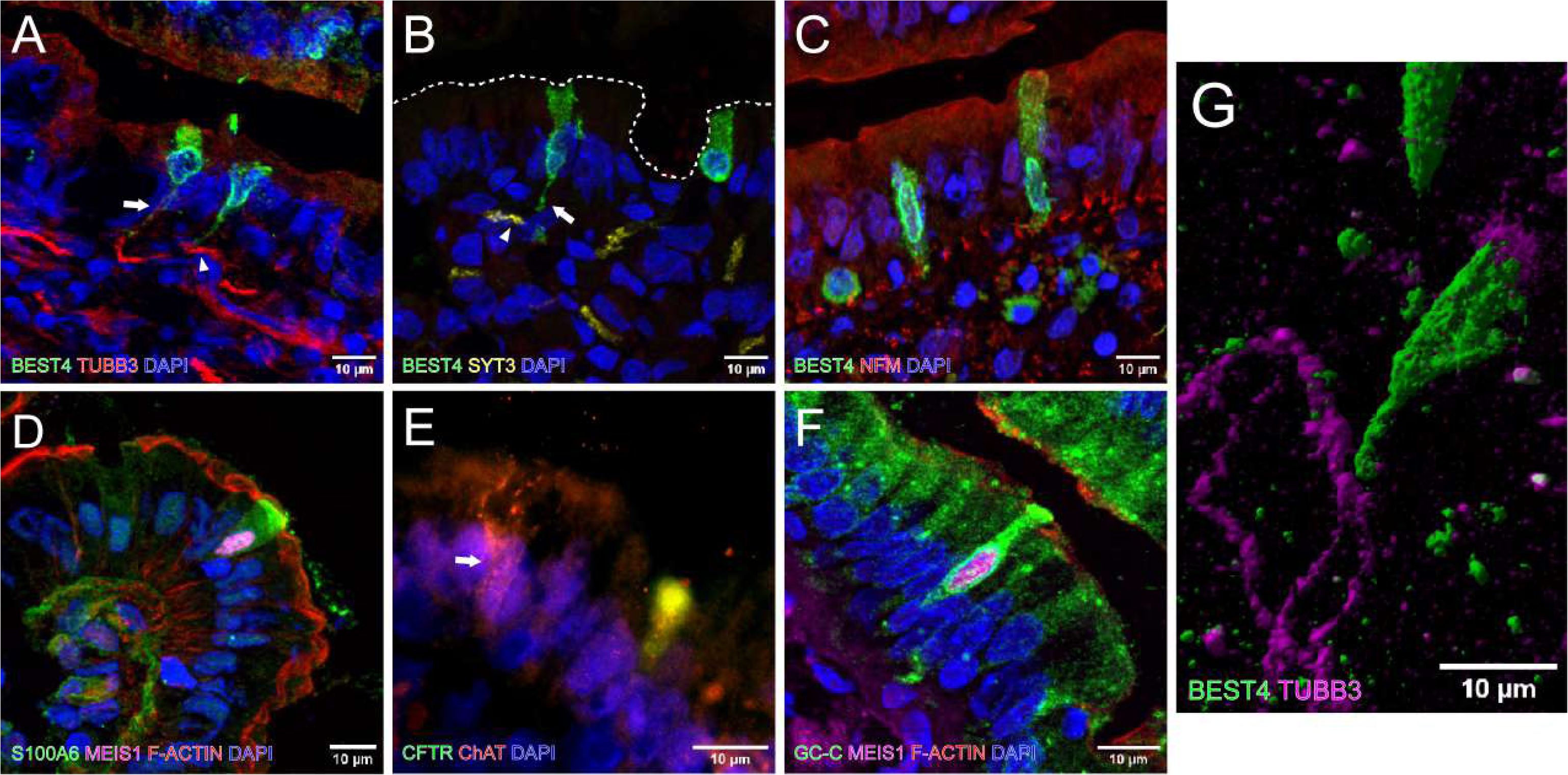
All regions are mid villus or villus tip rat epithelium. *(A)* CHE cells expressing BEST4 (green) have long basal pseudopodia *(arrow)* that extend to submucosal neurons *(arrowheads)* expressing TUBB3 (red). *(B)* A CHE cell basal pseudopod expressing BEST4 (green) extends to a submucosal neuron expressing SYT3 (yellow). *(C)* BEST4+ (green) CHE cells and all villus epithelial cells express NFM (red) at the basolateral membrane. *(D)* A MEIS1+ (magenta) CHE cell highly expresses cytoplasmic and apical S100A6 (green). *(E)* A CHE cell is innervated by neurons expressing CHAT (red). A goblet cell *(arrow)* expressing CHAT (red) is found in close proximity to the CHE cell. *(F)* A MEIS1+ (magenta) CHE cell highly expresses cytoplasmic and apical GC-C (green). GC-C is also in scattered in vesicular structures in villus enterocytes *(G)* BEST4 (green) and TUBB3 (magenta) staining shows CHEs at the same plane and close proximity to enteric neurons. Scale bars: 10 µm.

### CHE cells express neuropod genes and localize near submucosal neurons

Our previous studies on CHEs focused on CFTR localization and its apical trafficking into the brush border (6, 14, 34). ScRNA-seq data from rat jejunum allowed further investigations into other genes and proteins enriched and specific to CHEs, such as UGN and the basolateral channel BEST4, that enabled us to document that CHE cells possess basal processes that extend into the LP (Fig 2*A*, *B* and *G*). This observation coupled with the morphologic and phenotypic features of the novel intestinal neuropod cell (35) prompted further examination of the transcriptional profile from rat jejunum scRNA-seq data to identify CHE-enriched genes implicated in neuropod cell functionality. Indeed, neuropod genes were identified in human BEST4+ cells in the small intestine (9) and human CHEs (13). Neuropod cells are a newly identified type of sensory epithelial cell that forms synapses and transduce sensory signals from the intestinal lumen to the brain by fast neurotransmission onto neurons and the enteric nervous system (ENS) (36–38).

Neuropod-specific genes include genes involved in synaptic vesicle transmission and exocytosis, G-protein coupled receptors (GPCRs) and neuropeptides (35). IF labeling of tubulin beta 3 (TUBB3), a tubulin protein vital to microtubule formation in neurons and marker of myenteric neurons, and BEST4 revealed the proximity of CHE BEST4+ basal pseudopodia and TUBB3+ submucosal neurons (Fig 2*A*). 3D reconstruction of CHE-neuron proximity in thicker sections revealed that CHE basal pseudopodia are on the same plane as TUBB3+ neurons (Fig 2*G*).

ScRNA-seq data also identified synaptotagmin 3 (*Syt3*) transcription in CHEs, an important regulatory protein for synaptic activity (39). IF labeling of SYT3 and BEST4 visualized BEST4+ basal pseudopodia near SYT3+ neurons (Fig 2*B*). BEST4 is localized in the cytoplasm of CHE cells and pseudopodia that extend into the LP of the villus consistent with the morphology of neuropod cells (35, 37). The LP is innervated by submucosal neurons, providing neuropod cells a means to communicate with the ENS.

Neurofilament medium (NFM) is another neuronal protein implicated in intracellular transport to the axons and dendrites of neurons. BEST4+ CHE cells and indeed all villus epithelial cells were immunoreactive for NFM in the basolateral membrane, with significant fluorescence label in the membranes directly facing the LP (Fig 2*C*). CHE cells are also enriched for S100A6 with high protein concentration in the apical domain (Fig 2*D*). S100A6 is implicated in neuropod cell functionality and is linked to Ca^2+^-dependent insulin release, cell proliferation, vesicular transport and motility (40). Choline acetyltransferase (ChAT) synthesizes the neurotransmitter acetylcholine and is concentrated in the nerve terminal. Neuronal fibers expressing ChAT were found to innervate CHE cells on the villi near ChAT+ goblet cells, supporting acetylcholine-modulated mucus release (Fig 2E). CHE cells are often distributed in proximity to goblet cells throughout the villus, supporting a role for CHEs in mucus hydration and alkalinization. Lastly, the GC-C receptor, demonstrated to play a key role in regulating visceral pain, was recently shown to be a marker for neuropod cells in mice and human proximal small intestine (33). GC-C is also enriched throughout the cytoplasm of MEIS1+ CHE cells (Fig 2*F*).

### Uroguanylin is found ubiquitously in the rat intestinal epithelium and highly expressed in specific epithelial cells

UGN is distributed in the rat and human small intestine and colon in the apical membrane of all epithelial cells (28, 30). It is most abundant in the duodenum and lowest in the colon, consistent with its role as an integral regulator of CFTR-activated fluid and bicarbonate release where pH levels are lowest due to the influx of acidic content entering from the stomach (27). We recapitulated UGN protein distribution along the rostrocaudal axis of the rat intestine, spanning from the duodenum, jejunum, ileum, and colon (Fig 3*A*-*D*). In addition to its predominant distribution on the apical domain of villus epithelial cells, we found UGN-rich cells interspersed on the villi (Fig 3*A-D*). IF labeling of CFTR and UGN revealed that some CHE cells also express very high levels of UGN in the subapical compartment and apical domain (Fig 3*E*). Consistent with previous published findings (28), we found some UGN+ cells were also enriched for the EEC marker CHGA (Fig 3*F*). Interestingly however, not all UGN+ cells were CHEs, and not all UGN+ cells express CHGA.

**Fig 3.**
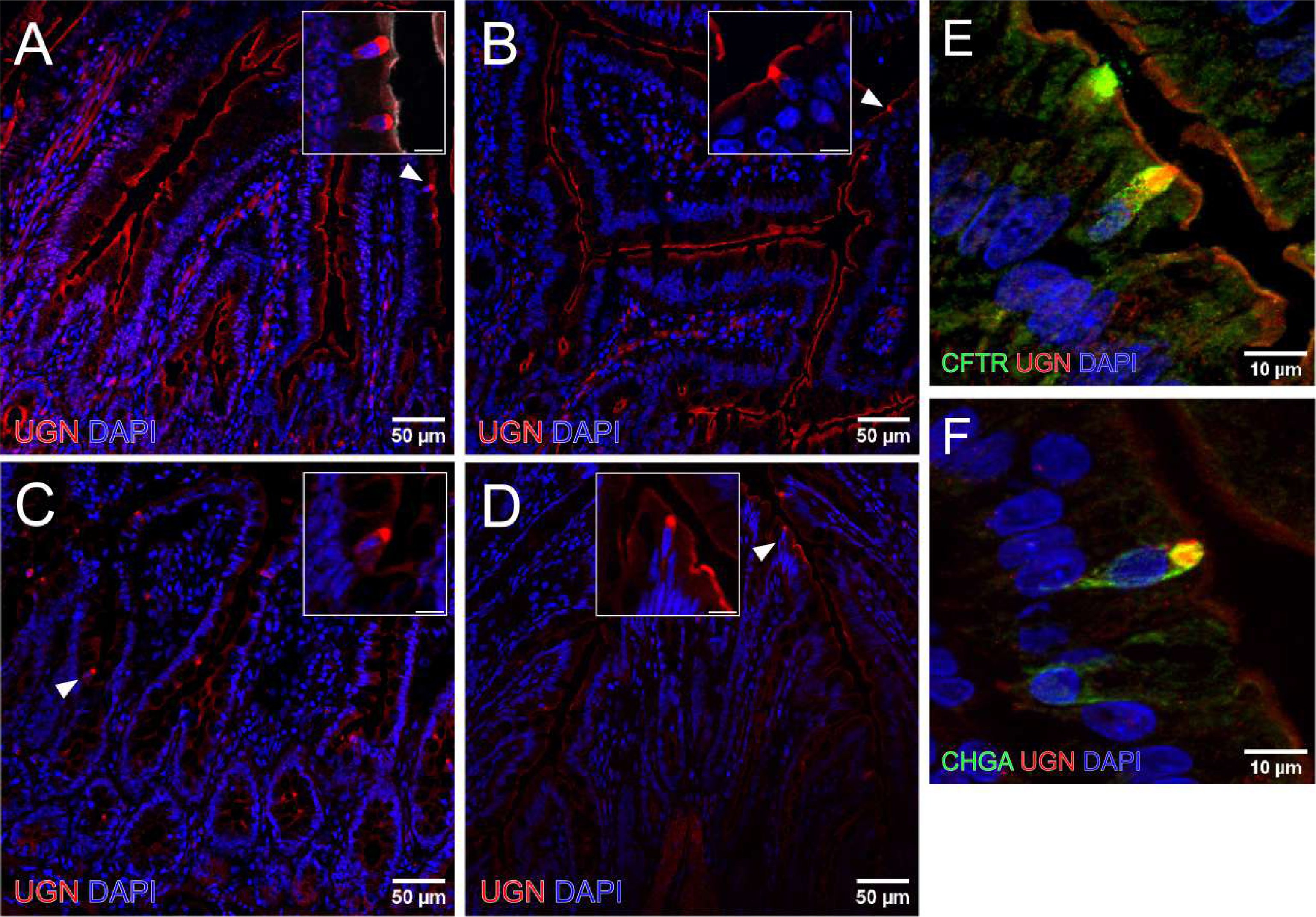
Rostrocaudal axis of rat intestine immunolabeled for UGN (red): *(A)* duodenum, *(B)* jejunum, *(C)* ileum, and *(D)* colon. UGN is localized on the apical membrane of all epithelial cells, and UGN fluorescence label decreases along the axis. *(A-D)* Specific epithelial cells *(arrowheads)* highly express UGN in the apical region. *(E)* Some cells with abundant UGN (red) also highly express CFTR (green). *(F)* Another subpopulation of epithelial cells highly expressing UGN (red) also express CHGA (green). Scale bars: *(A-D)* 50 µm, *(inserts)* 10 µm, *(E* and *F)* 10 µm.

### CHE cells sense and respond to low pH by rapidly trafficking CFTR, GC-C and OTOP2 to the apical domain

ScRNA-seq identified high levels of the acid-sensing GPCR receptor *Gpr4* and *Gpr6* in rat CHE cells (Fig 1*A*), indicating a functional role for CHEs in sensing pH. The lack of effective working antibodies made it difficult to demonstrate GPR4 immunoreactivity in CHEs in rat tissues. Activation of GC-C and the cGMP pathway in epithelial cells within acidic regions of the intestine stimulates HCO_3_^-^ secretion via CFTR (27), prompting us to investigate whether luminal exposure to low pH alters the subcellular distribution of CFTR and GC-C within CHE cells in the proximal jejunum.

In control tissue treated with saline, CFTR and GC-C were observed in both the apical and subapical domain of all CHEs (Fig 4*A* and *C*). The fluorescence intensity ratio of CFTR and GC-C comparing the apical and subapical region revealed a mean 3.6-fold and 1.5-fold increase in fluorescence per CHE cell, respectively, following exposure to low pH consistent with trafficking activity (Fig 4*E* and *F*). Following exposure to a low pH solution for 30 min, CFTR and GC-C trafficked significantly to the brush border membrane (BBM) of CHEs in both crypts and villi, consistent with previous findings (Fig 4*B* and *D*) (15). The CFTR apical to subapical fluorescence ratio increased 4.9-fold and GC-C increased 2.4-fold (Fig 4*E* and *F*). We observed no change in the number of CHE cells following luminal exposure to low pH (Fig 4*G*). Additionally, the acidic pH introduced into the lumen was effectively and rapidly neutralized (Fig 4*H*). Successful buffering of the acidic luminal contents, despite no significant difference in CHE abundance, suggests that CFTR and GC-C activity in CHE cells play a key role. There was no change in HCO_3_^-^ *(I)* and Cl^-^ *(J)* (mmol/L) concentrations across treatment and control groups. Collectively, these findings support their critical function in sensing and regulating pH by driving CFTR and GC-C trafficking.

**Fig 4.**
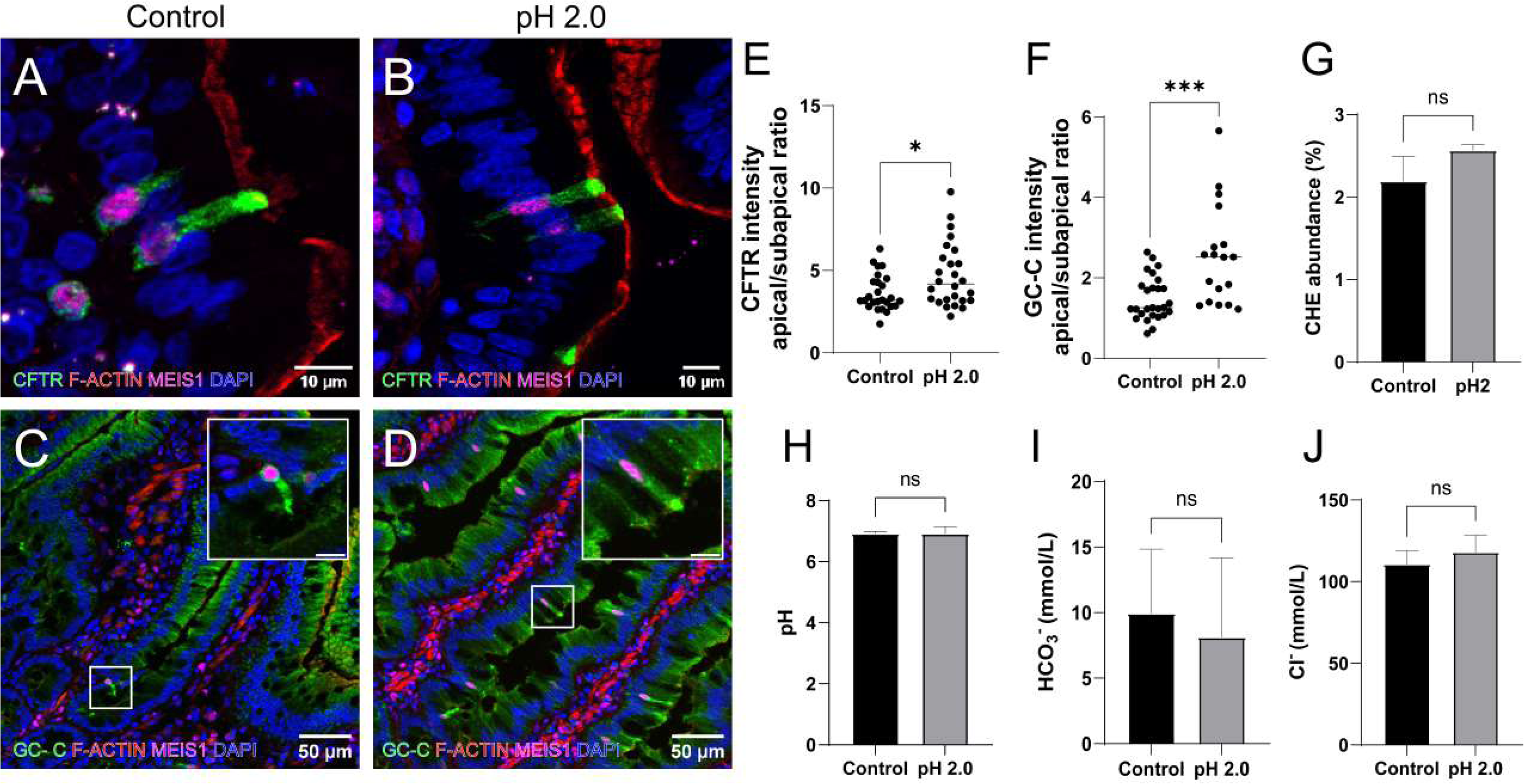
Normal rat jejunum treated with *(left)* buffered saline (pH 7) or *(right)* pH 2 solution for 30 min immunolabeled for *(A* and *B)* CFTR (green), MEIS1 (magenta), F-actin (red), and *(C* and *D)* GC-C (green), MEIS1 (magenta), F-actin (red). *(B*) Apical CFTR fluorescence is increased in MEIS1+ CHE cells in low pH-treated animals. *(D*) GC-C expression is significantly greater in both MEIS1+ CHE cells and the villus epithelium. (*E)* There was no significant change in the abundance of CHE cells (percent of all epithelial cells) after low pH exposure. *(F* and *G)* Apical to subapical CFTR and GC-C fluorescence intensity ratio significantly increased following pH 2 treatment. *(H)* The pH of luminal fluids collected from saline control and pH 2 animals was not affected. *(I* and *J)* There was no change in HCO_3_^-^ and Cl^-^ (mmol/L) concentrations across treatment groups. P-values: *<0.05, **<0.001. Scale bars: *(A* and *B)* 10µm, *(C* and *D)* 20 µm.

CHEs express CFTR, GPR4, GC-C, UGN, BEST4 and OTOP2, all involving in sensing and responding to acidic pH. We hypothesized that the proton channel OTOP2 may undergo changes in expression in response to low luminal pH *in vivo.* Immunolabeling of OTOP2 and MEIS1 confirmed high protein expression in CHE cells but only when exposed to low pH (Fig 5*B*). Immunolabel for OTOP2 protein was not detected in CHEs in normal rat jejunum (Fig 5*A*). However, it dramatically increased at the apical domain of MEIS1+ CHE cells after 30 min exposure to a low pH environment. The rapid apical recruitment and accumulation of OTOP2 are consistent with CHEs sensing and responding to changes in luminal pH.

**Fig 5.**
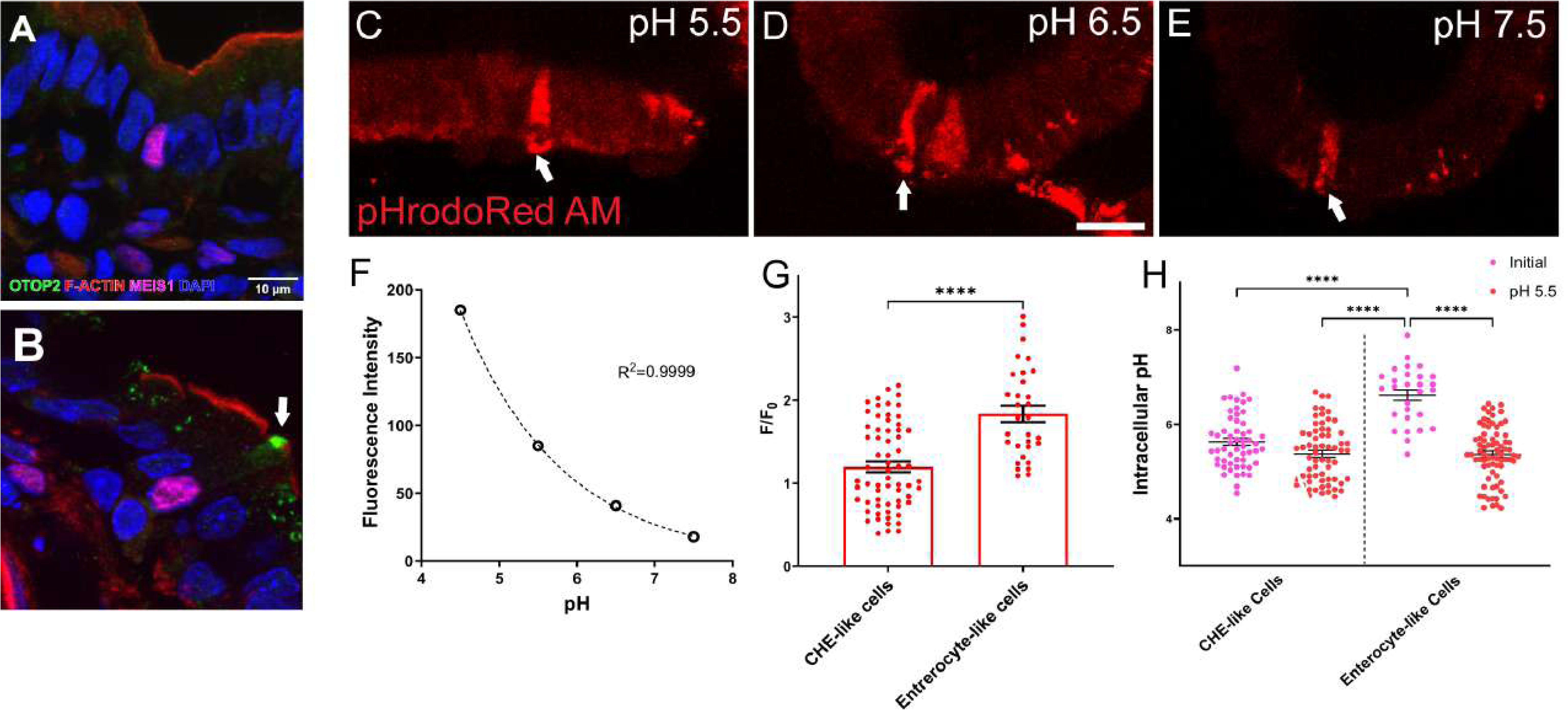
Normal rat jejunum exposed to *(A)* buffered saline (pH 7) or *(B)* pH 2 solution immunolabeled for OTOP2 (green), MEIS1 (magenta) and F-actin filaments (red). *(A)* Saline-treated control rats showed no detectable OTOP2 expression in CHE or other villus epithelial cells. *(B)* Rats treated with pH 2 solution demonstrated significant upregulation of OTOP2 in the apical membrane of MEIS1+ CHE cells (arrow). Organoids incubated in pH 5.5 buffer *(C)*, pH 6.5 buffer *(D)*, and pH 7.5 buffer *(E)*. High pHrodo Red AM fluorescence indicates the presence of CHE-like cells with lower intracellular pH (arrow). Scale bar: 20 μm. (F) Calibration curve showing pHrodo Red AM fluorescence intensity in response to pH buffers 4.5, 5.5, 6.5, and 7.5. *(G)* Fluorescence ratio (F/F) in organoids at pH 5.5 normalized to initial pH 7.5. *(F)* Intracellular pH comparison based on the calibration curve (F) for organoids in initial pH 7.5 and pH 5.5 buffers. Data in *(G)* and *(H)* represent mean ± SEM, with Mann-Whitney test applied and *p* ≤ 0.0001.

In human colon, BEST4+/OTOP2+/CFTR- cells were found to sense and transport protons into the cell in response to lowering the extracellular pH (41). Akin to BEST4+ cells in the colon (9, 41), functional studies using pHrodo Red AM, a membrane-permeable pH indicator dye, showed that CHE-like cells in the small intestine can be responsive to changes in extracellular pH, much like BEST4+ cells in the human colon. In rat jejunal organoids exposed to different pH conditions, a global increase in pHrodo Red AM fluorescence was observed as the extracellular environment became more acidic. (Fig. 5*C-H*). However, a subset of cells with a morphology resembling CHE cells (CHE-like cells) exhibited a more pronounced increase in fluorescence compared to enterocyte-like cells, particularly between pH 5.5 and pH 6.5 buffers (Fig. 5*C-E*, *G* and *H*). This suggests that CHE-like cells may have slightly increased sensitivity to extracellular pH changes relative to other epithelial cell types, due the expression and activity of proteins as BEST4 and OTOP2.

Thereby, BEST4+ cells in the small intestine (CHEs) and colon can respond to low pH, but CFTR is only found in BEST4+ cells in the proximal small intestine. Our observations that CFTR is absent from the apical domain and brush border of CHEs in ΔF508 CHE cells (Fig. 9*D* and *E*) coupled with knowledge that the CF intestine is more acidic than normal suggest that CHEs play a critical role in CF pathogenesis in the intestine by sensing and responding to low pH. Whether OTOP2 and other acid receptors in CHEs are dysregulated in CF warrants investigation.

### CHE cells sense and respond to STa by rapidly trafficking CFTR and GC-C to the apical membrane

STa and its endogenous peptides, guanylin and UGN, bind to the GC-C receptor and activate its intracellular catalytic domain, resulting in the hydrolysis of guanosine triphosphate (GTP) and accumulation of intracellular cyclic GMP (cGMP) levels (42, 43). We demonstrated previously that cGMP signals cGMP-dependent protein kinase (cGKII) to phosphorylate CFTR resulting in its phosphorylation and trafficking to the BBM of enterocytes, leading to an efflux of Cl^-^ and water into the lumen (20, 44). Since CHEs are enriched with GC-C and the cGMP machinery to activate CFTR, we examined whether luminal exposure to STa alters the subcellular distribution of CFTR, GC-C and BEST4 in CHEs in the proximal jejunum. After 30 minutes of luminal stimulation with STa, CFTR (Fig 6*A*, *B* and *G*) and GC-C (Fig.6*E*, *F* and *H*) trafficked to the BBM of CHEs and enterocytes. Not surprisingly, STa, a cGMP agonist, did not elicit changes in BEST4 localization in CHEs, consistent with its activation by calcium (Fig 6*C* and *D*). CHE abundance did increase slightly after STa treatment (Fig 6*I*).

**Fig 6.**
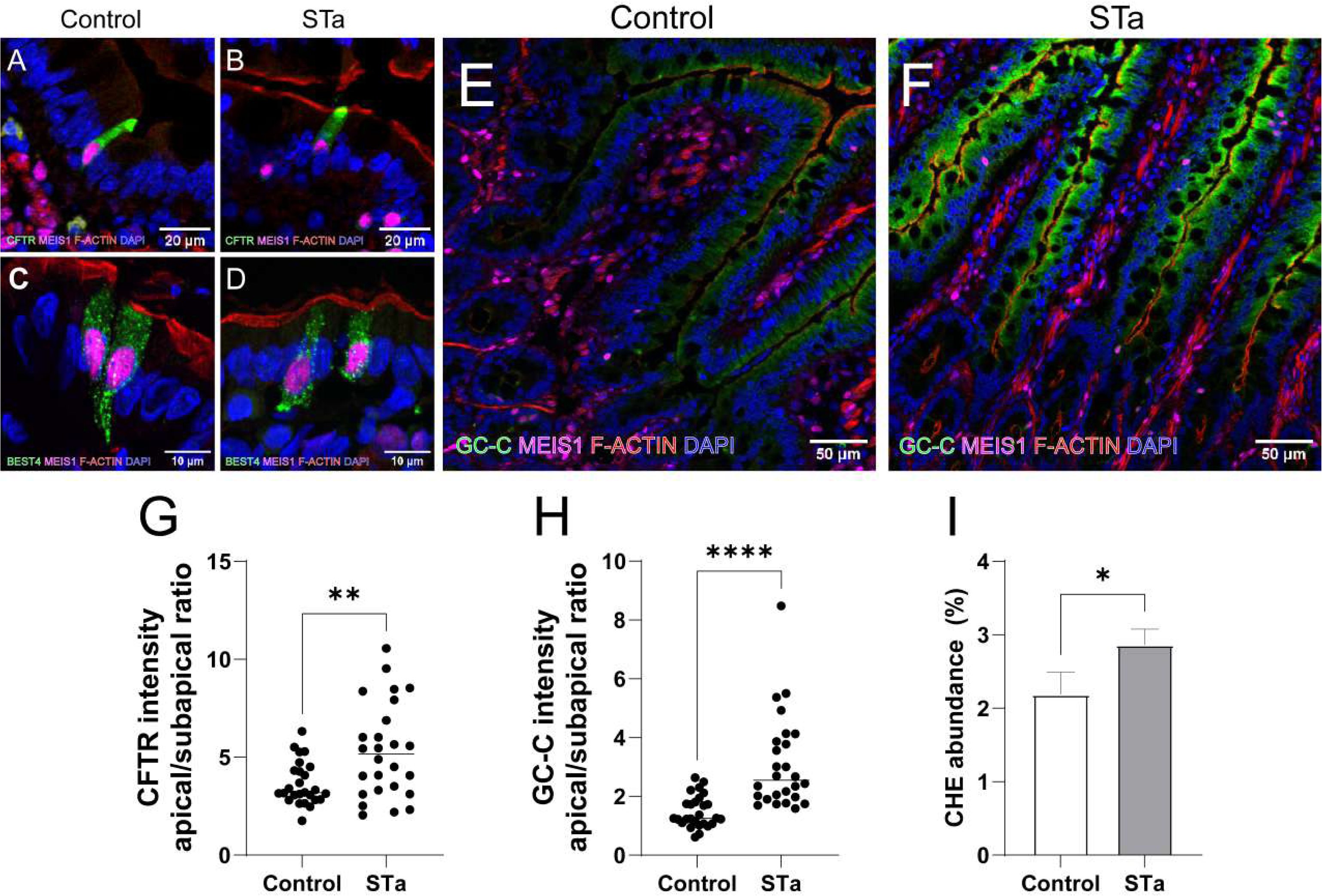
Normal rat jejunum treated with *(left)* buffered saline or *(right)* STa for 30 min. All tissues immunolabeled for MEIS1 (magenta), F-actin (red), and *(A* and *B)* CFTR (green), *(C* and *D)* BEST4 (green), or *(E* and *F)* GC-C (green). *(A* and *B)* Fluorescence imaging of CFTR indicates increased concentration in the apical membrane of MEIS1+ CHE cells in STa treated animals *(B)*. *(G)* Apical to subapical ratio quantification of CFTR fluorescence intensity reveals increased trafficking to the apical membrane in CHE cells after STa treatment. *(C* and *D)* BEST4 fluorescence suggests no significant change in protein abundance or localization in CHEs after STa treatment. *(E* and *F)* GC-C levels are heightened across the jejunum in STa-treated animals. *(H)* Apical to subapical ratio quantification of GC-C fluorescence intensity shows increased trafficking to the apical membrane of MEIS1+ CHE cells. *(I)* CHE abundance (percentage of all epithelial cells) is not altered between saline and STa-treated rats. Scale bars: *(A-D)* 10 µm, *(E* and *F)* 50 µm.

### MYO1B traffics CFTR to the apical membrane in CHE cells

We suspected MYO1B to be the motor protein responsible for trafficking CFTR to the apical membrane of CHE cells from subapical vesicles. As reported previously, MYO1A has been shown to traffic CFTR to the brush border membrane of villus enterocytes (26). This specific molecular motor is not present in CHEs, suggesting that CHE cells are unique in their regulation of CFTR trafficking (6). Following IF confirmation of CFTR and MYO1B co-localization in CHEs (Fig. 1*E* and *F*), we further investigated MYO1B-dependent trafficking of CFTR by generating a MYO1B knockdown (KD) rat intestinal organoid model (21) (Fig 7*A-D*).

**Fig 7.**
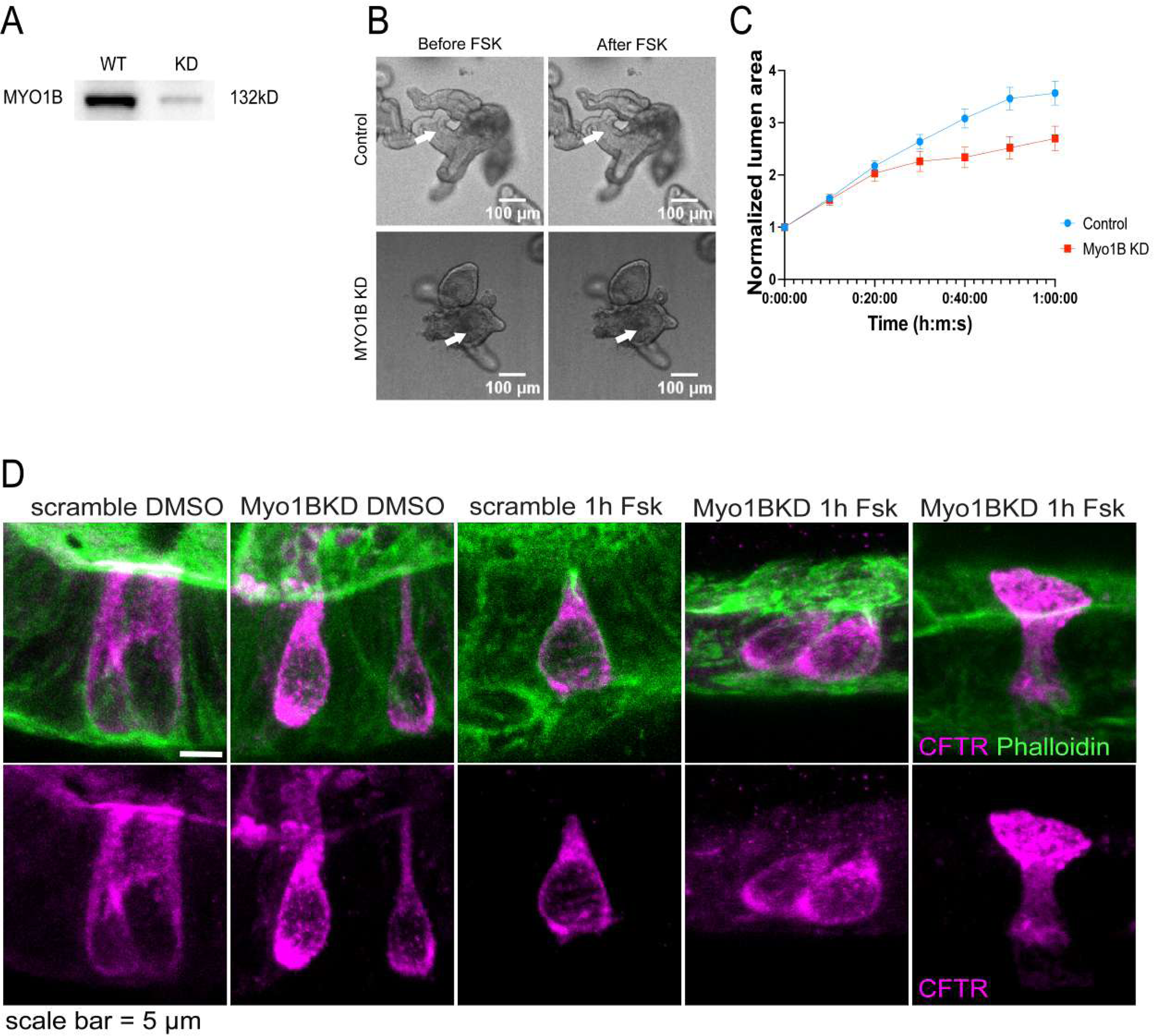
(A) Protein concentration analysis confirms successful knockdown of MYO1B via siRNA (70% efficiency). *(B)* Normal and MYO1B knockdown (KO) organoids were treated with dimethylsulfoxide (DMSO) or forskolin (FSK) for 1 hour (h), respectively, and lumen swollen area was quantified in *(C)*. Scale bars: *(B)* 100 µm. *(D)* Normal and MYO1B KO organoids treated with DMSO or FSK for 1 h were immunolabeled for CFTR (magenta) and phalloidin (F-actin filaments, green). There were no significant changes in organoid swelling after FSK stimulation in MYO1B knockdown KO organoids, but CFTR apical immunolabel appeared more diffuse. Scale bars: *(D)* 5 µm.

Although small hairpin RNA (shRNA)-mediated knockdown of MYO1B (Fig 7*A*) in rat intestinal organoids did not result in a significant reduction in CFTR trafficking in CHE cells in vitro (Fig 7*D*), this suggests that similar to our observations in Myo1a/Myo6 double KO mice (17), alternate plus end myosins including Myo 1c, Myo1d, Myo1e, Myo5a may compensate for the loss of MYO1B function (26).

### Expression of *Cftr* and G*uca2b* transcripts in ΔF508 rat CHEs

Nothing is known about the role of CHEs in the CF intestine. To identify changes in disease-relevant mRNA transcripts in CHEs in CF, we examined *Cftr* and *Guca2b* mRNA abundance and localization in wild-type (Fig 8*A-D* and *I-L*) and ΔF508 (Fig 8*E-H* and *M-P*) rats. FISH using a CFTR probe revealed loss of *Cftr* mRNA abundance in ΔF508 CHE cells (Fig 8*E* and *F*) and along the rostrocaudal intestine (Fig 8*M-P*). Reduction of the quantity of *Cftr* mRNA in CHE cells in ΔF508 rats is consistent with this class II mutation in the *CFTR* gene in CF (45). However, there was no change in the localization of *Guca2b* in CHE cells (Fig 8*G* and *H*) or along the intestine (Fig 8*M-P*), suggesting that these cells are still enriched with *Guca2b* transcript and *Guca2b* expression is not dependent on CFTR in CHE cells. We also observed *Guca2b* highly-expressing cells in the differentiated colon compartment of both wild-type and ΔF508 animals, that are likely BEST4+ colonic cells (Fig 8*L* and *P*).

**Fig 8.**
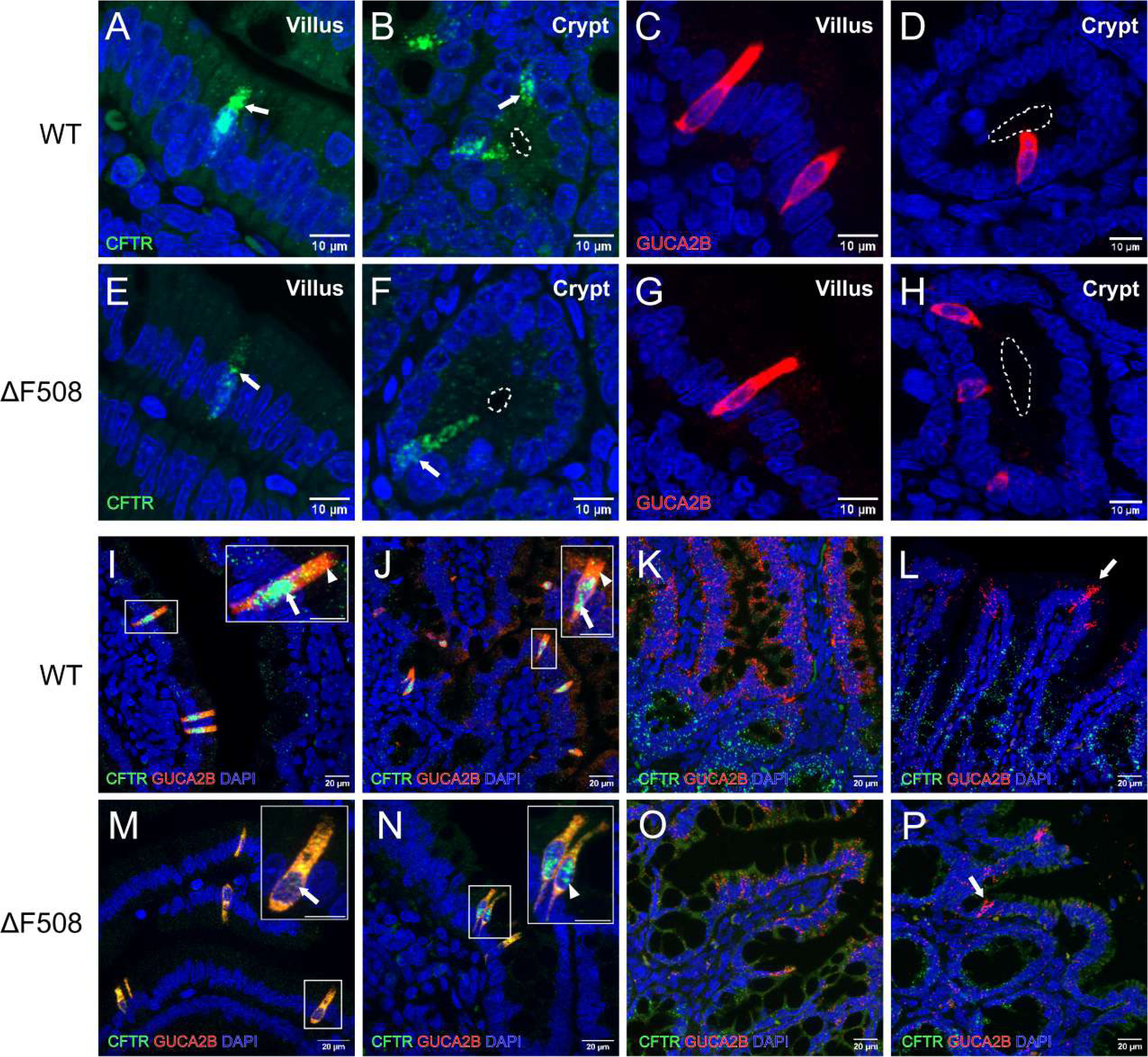
Significantly higher *Cftr* mRNA expression in CHEs *(arrow)* is observed in villi *(A)* and superficial crypts *(B)* of wild-type (WT) rats compared to other enterocytes and CHEs (*arrow*) present in the villi *(E)* and *(F)* superficial crypts of ΔF508 rats. Notably, there is also a reduction of *Cftr m*RNA in ΔF508 rat enterocytes *(F)*. Higher *Guca2b* mRNA expression is observed in scattered cells resembling CHEs in villi *(C)* and superficial crypts *(D)* of WT and in villi *(G)* and superficial crypts *(H)* of ΔF508 rats. Along the rostrocaudal axis of the small intestine *((I* and *M)* duodenum, *(J* and *N)* jejunum, and *(K* and *O)* ileum) and large *((L* and *P)* colon) intestines, tissue was double-labeled for *Cftr* (green) and *Guca2b* (red) mRNA in WT and ΔF508 rats, respectively. *Cftr* and *Guca2b* mRNA abundance suggest CHE cell transcriptional profile and morphology in WT rat duodenum *(I)* and *(J)* jejunum. In these cells, *Cftr* mRNA is predominantly localized in the nuclei *(arrow)*, while *Guca2b* mRNA is abundant in the cytoplasm. CHE cells were not observed in the ileum *(K)* or colon *(L)*. However, a rare population of cells with high *Guca2b* mRNA levels *(arrow)* is found in the colon of both WT *(L)* and *(P)* ΔF508 rats. *Cftr* mRNA abundance is significantly reduced in the intestine of ΔF508 rats *(M-P)*. Cells with high higher *Guca2b* mRNA levels in ΔF508 rat exhibited either complete *(M, arrow)* or partial *(N, arrowhead)* loss of *Cftr* mRNA in the nuclei. *Cftr* mRNA is still detected in the crypts of ΔF508 rats (*F*, *N*, *O*, and *P*) but at lower levels. *Guca2b* mRNA levels are not affected by loss of *Cftr*. Scale bars: *(A-H)* 10 µm, *(I-P)* 20 µm.

### CHE cells differentially express CFTR and GC-C protein in ΔF508 and CFKO rat jejunum

Remarkably, the distribution of CHEs and expression of CFTR and CHE-specific proteins in the CF intestine remain unknown. The absence of CHEs in mice, lack of access to CF human proximal small intestine and working antibodies against human CFTR have contributed to the ongoing challenges contributing to lack of progress in this area. However, successful generation of ΔF508 and CF-knockout (CFKO) rat models provides a novel opportunity to characterize CHE cells and proteins of interest in CF intestinal disease. These rat CF models are of particular importance to understand the role of CHEs in CF. We first examined the morphology of normal, ΔF508 and CFKO rat intestine in hematoxylin and eosin (H&E) stained sections (Fig 9*A-B*). Consistent with CF human disease, we observed increased lymphocyte infiltration in the lamina propria of the ΔF508 animal (Fig 9*B*), and increased abundance of goblet cells in CFKO intestine (Fig 9*C*).

**Fig 9.**
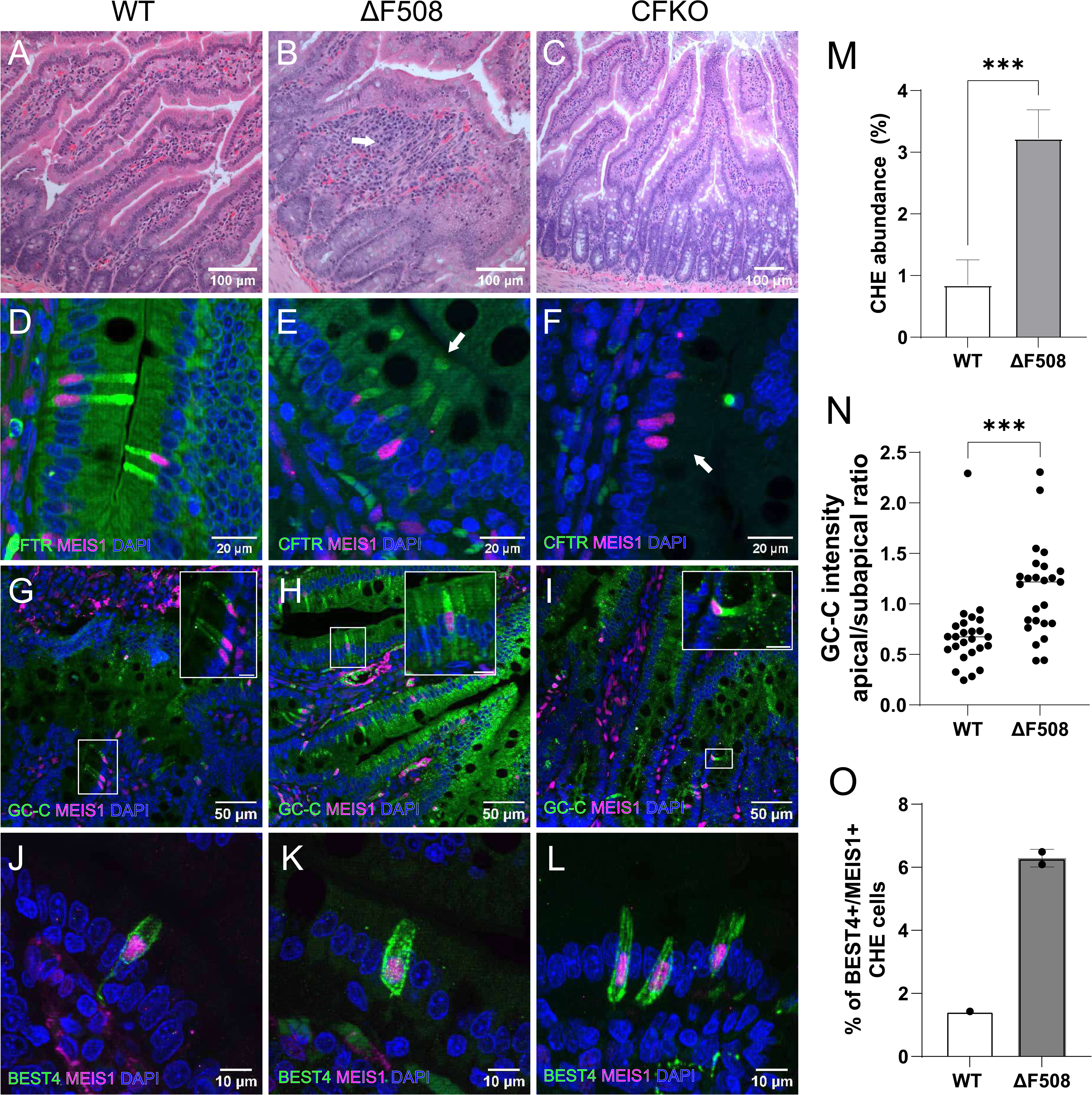
All tissues shown are rat jejunum. CF conditions are organized accordingly: *(left)* normal (wild-type (WT)), *(center)* ΔF508, *(right)* CFKO. *(A-C)* Hematoxylin and eosin (H&E) staining of crypt-villus axis. *(B)* Increased abundance of lymphocytes in villus submucosa *(arrow)* is consistent with human CF intestinal disease. *(D-F)* Villus epithelium immunolabeled for CFTR (green) and MEIS1 (magenta). ΔF508 rats show loss of CFTR protein *(E)* in the apical domain and membrane of CHE cells (*arrow*) and markedly reduced CFTR in the cytoplasm. CFKO rats demonstrate total absence of CFTR *(F)* in the entire villus epithelium. Both ΔF508 and CFKO animals retain MEIS1+ nuclei in CHE cells (*E* and *F*). *(G-I)* GC-C (green) and MEIS1 (magenta) immunolabel in WT *(G),* ΔF508 *(H)*, and CFKO *(I)* rats. GC-C is higher in villus enterocytes of ΔF508 rat jejunum (Fig 9*H*) and was preserved in MEIS1+ CHE cells in CFKO (Fig 9*I*). *(J-L)* BEST4 expression in CHE cells does not change in ΔF508 and CFKO animals. Scale bars: *(A-C)* 100 µm, *(D-F, J-L)* 10 µm, *(G-I)* 50 µm. *(M)* A 3-fold increase in CHE abundance is observed in ΔF508 compared to WT rats. *(N)* GC-C apical/subapical ration intensity increases in ΔF508 compared to WT rats. *(O)* BEST4+MEIS+ cell abundance in WT and ΔF508.

ΔF508 CFTR is the most common disease-causing mutation in CF patients. ΔF508 is a Class II mutation that prevents the proper folding, intracellular apical traffic and function of the protein. Consistent with the ΔF508 trafficking defect, the ΔF508 rat displayed marked decreased CFTR protein in MEIS1+ CHEs and other villus epithelial cells, in addition to a specific loss of CFTR in the apical domain and brush border (Fig 9*E*). In the CFKO rat, complete loss of CFTR was observed in the entire villus epithelium (Fig 9*F*). Interestingly, MEIS1+CHE cells presented a 3-fold abundance (Fig 9*M*) in the ΔF508 CF intestine compared to WT rat. This suggests that fate specification and differentiation of CHEs is independent of CFTR expression.

Since we identified high levels of GC-C in CHEs, and the cGMP machinery and pathway are crucial to acid-stimulated activation of CFTR and HCO_3_^-^ secretion in the proximal small intestine, we examined GC-C distribution in ΔF508 and CFKO rat jejunum to determine whether CF intestinal disease is associated with changes in GC-C. GC-C immunolabel was higher in villus enterocytes of ΔF508 rat jejunum (Fig 9*H*) and was preserved in MEIS1+ CHE cells in CFKO (Fig 9*I*). Again, it is noteworthy that MEIS1+ CHE cells express GC-C despite the loss of CFTR. CHE cell differentiation and expression of MEIS1 and GC-C are irrespective of CFTR activity, suggesting a complex CHE fate. Consistent with our observations on GC-C, BEST4 localization in MEIS1+ CHE cells is preserved in ΔF508 and CFKO rat intestine (Fig 9*K* and *L*). BEST4 labels MEIS1+ CHE cells in the basolateral domain across all three conditions, including in CFKO, indicating that BEST4 is preserved in the absence of CFTR.

## DISCUSSION

CHEs were first identified in 1995 by immunolocalization based on the unusually high levels of CFTR protein in the cells (3). Our laboratory published several papers characterizing the ultrastructure, subcellular localization of CFTR, and the behavior of CFTR, ion channels, molecular motors and proteins identified in CHEs under various conditions *in vivo* in rat intestine (3, 4, 6, 14). However, several limitations precluded investigations to gain further insight into this intriguing subpopulation of cells; including the lack of specific markers to facilitate enrichment of CHEs, the absence of CHEs in mice, and insufficient tools to identify genes specifically enriched in CHEs. ScRNA-seq studies in human duodenum provided the first window into CHE-specific transcriptome in human intestine that confirmed high BEST4 expression in CHEs, resulting in the designation BCHE (13). Several groups subsequently used scRNA-seq approaches to report the transcriptomic profiles of rare cell types in native intestinal tissues in a number of species, and disease conditions (9, 10, 41, 46, 47). In contrast to CHEs that are confined to the proximal small intestine of rats and humans, BEST4+ cells were identified throughout the intestine in several species excluding mice (8). CFTR transcript was identified only in BEST4+ cells in the proximal small intestine, but interestingly, the link between BEST4+/CFTR+ cells and CHEs was not firmly established. This is likely in part due to lack of data confirming the expression of BEST4 in CHEs in the rat proximal small intestine.

The current study used scRNA-seq data from rat jejunum to identify and examine disease-relevant genes specifically upregulated in CHEs. Using antibodies raised against these proteins, we examined the cellular and subcellular distribution of CHE-specific proteins in normal rat under physiologic and acidic luminal conditions consistent with CF intestinal disease. Since very little is known about the importance of CHEs in CF intestinal disease, we examined changes in disease-relevant CHE-specific proteins in CHEs in normal, ΔF508 CF and CFKO rats.

Like CHEs, rare cells in the lung called pulmonary ionocytes express very high levels of CFTR. FOXI1 is the major transcription factor regulating ionocytes (23, 24). We sought evidence for *Foxi1* expression in rat CHEs but failed to identify it in our scRNA-seq data. Instead, *Meis1* was identified as a transcription factor enriched and specific for CHE cells. Antibody label of sections from mouse trachea confirmed FOXI1 and CFTR protein enrichment in ionocytes, while rat jejunum showed an absence of FOXI1 in CHEs and intestinal cells. MEIS1 specificity for CHEs was further confirmed by immunostaining and used as a marker of CHEs in our double staining experiments to further characterize proteins enriched in CHEs. These findings confirm that while pulmonary ionocytes and CHEs both express high CFTR, their transcriptional and cell fate specification are different.

*Myo1b* transcript enrichment in CHEs was confirmed at the protein level by immunolabel of rat jejunum villus CHEs. MYO1A, the motor that is required for villus enterocyte brush border development, regulates CFTR traffic from subapical endosomes into the brush border of mouse intestine. Rat villus CHEs lack MYO1A (6) but robustly traffic cargo from subapical endosomes into the brush border membrane. ScRNA-seq and protein immunolabel identified enrichment of *Myo1b*/MYO1B in CHE cells, suggesting that MYO1B was the motor implicated in CFTR brush border trafficking in CHE cells. Surprisingly, efficient silencing of MYO1B in rat organoids reduced CFTR-activated fluid in CHE-expressing organoids but did not eliminate CFTR traffic into the brush border of CHEs. This observation is consistent with our previous observations of compensatory myosins in transgenic mouse models. MYO1A-MYO 6 double KO mice generate alternate myosins to traffic cargo in the absence of MYO1A and MYO6 (17). IF label also confirmed tubulin 2 enrichment in CHEs, consistent with ultrastructural evidence of prominent motor fibers traversing the cells (4).

*Guca2b* and *Guca2a* transcript enrichment in BEST4+ cells in the intestine were identified in several studies. We confirmed that BEST4 protein is indeed enriched in the basal domain of CHEs, localized to punctate vesicular like structures and extends into basal process. Previous studies focused on apical traffic of CFTR in CHEs and thus failed to identify basal processes extending to the LP. *GUCA2B* encodes for the hormone UGN, a key regulator of cGMP and acid-stimulated HCO_3_^-^ secretion via CFTR. Unlike *GUCA2A* that encodes guanylin, that is more abundant in the distal alkaline segments of intestine, *GUCA2B* is distributed more abundantly in the acidic proximal small intestine. Validated UGN antibodies allowed confirmation that CHEs indeed express high levels of UGN in the cytoplasm that extends into its basal process. FISH studies also confirmed high levels of *Guca2b* co-localizing with *Cftr* mRNA in CHEs. UGN and GC-C enrichment in CHEs align with a prominent role for CHEs and CFTR in regulating HCO_3_^-^ secretion to buffer the acidic effluent from the stomach entering into the proximal small intestine. Consistent with this, CFTR, GC-C and the proton channel OTOP2 were highly upregulated in the apical domain of CHEs under acidic conditions. BEST4+ cells in the colon also express UGN and OTOP2 but lack CFTR. Functional studies confirmed proton uptake under acidic conditions, likely by setting cGMP tone in the colon. Our data suggest that CHEs are potential sensors of luminal pH in the proximal small intestine, and the role of CFTR in CHEs appears to be critical for responding to an acidic luminal pH, by secreting HCO_3_^-^ into the lumen. This is further supported by the finding that CFTR is absent in the apical domain of CHEs and intestinal cells in ΔF508 CF disease whose hallmark is an acidic lumen. We confirmed the predicted GC-C localization on the apical domain of villus enterocytes, but its localization and enrichment in vesicles within the cytoplasm of CHEs was surprising. Apical traffic of GC-C in response to an acidic lumen and its apical enrichment in CHEs in ΔF508 CF suggests that GC-C is central to acid-stimulated signaling under physiologic conditions and in CF.

Neuropod cells are sensory epithelial cells that form synapses onto neurons to transduce sensory signals from the intestinal milieu to the brain, using neurotransmitters. CHEs fit the criteria for neuropod cells as they display the morphologic features with basal processes, proximity to neurons, and express the molecular GPCR acid sensor, GPR4, and the neuronal sensor SYT3. Recent studies identified GC-C-rich neuropod cells in mouse and human small intestine and confirmed a critical role for GC-C in reducing visceral pain (VP) upon stimulation with cGMP agonists. Interestingly, GC-C-rich neuropod cells in mice and human express markers of EECs and neurons but lack hormones or machinery to regulate electrolyte transport. CHEs differ from GC-C-rich neuropod cells. Although CHEs are enriched for GC-C, express neurotransmitters, and are in close proximity to neurons, they are not present in mice, but are enriched for the hormones *Guca2a* and *Guca2b*, and express cellular machinery to regulate ion transport including CFTR, NKCC1 and BEST4. Why BEST4 and UGN are in basal processes and the role of GC-C in CHEs require further investigations.

The enterotoxin STa, a potent activator of cGMP-stimulated CFTR-mediated fluid secretion in the small intestine, leads to diarrhea. Previous studies in rat jejunum examined traffic of CFTR onto the surface of enterocytes (20) in STa-stimulated fluid secretion, but did not examine the impact of STa on CHEs or GC-C. The finding here that STa-stimulated apical traffic of CFTR in CHEs and GC-C in villus enterocytes was not surprising, given the abundance and distribution of GC-C in punctate structures in CHEs and in villus enterocytes. But it provides new evidence of GC-C regulation by traffic in native tissues.

High expression of CFTR in ionocytes suggested that they could be a major site of chloride secretion and target in CF lung disease, but functional studies indicated that they are not a major site of secretion (48), but instead mediate chloride absorption (49). More recent studies indicate that ionocytes regulate airway surface liquid (ASL) pH by secreting bicarbonate (50). Although CHEs and ionocytes share a similar rare distribution and high CFTR expression, their transcriptomic profiles are quite different. For example, BEST4, a major marker of CHEs on the basal membrane, is not expressed in ionocytes that instead express barttin chloride channels (49). Furthermore, ionocyte abundance was unchanged in CF (51), but was decreased in nasal mucosa of children with chronic rhinosinusitis (52). Although a secretory function for CHEs has not been confirmed, they express abundant hormones, ion channels and signaling machinery that regulate electrolyte secretion. Furthermore, CHEs lack absorptive markers, robustly traffic CFTR and GC-C upon activation of secretion and exposure to low pH, and human CHEs arise from secretory progenitors. In contrast to human ionocytes, CHE abundance is higher in the CF rat intestine, and GC-C is upregulated while BEST4 and MEIS1 are preserved.

Our analyses of CHE cells have identified: (1) a capacity for sensing and regulating pH in the proximal small intestine; (2) their connection to the enteric nervous system (ENS) that mediates rapid responses to changing luminal conditions, including STa; and (3) changes in CHE-specific protein distribution that have significant implications for intestinal CF pathogenesis. *In vitro* assessment of CHEs further elucidated the machinery involved in CFTR trafficking to the brush border, playing a critical role in CHE function. Our characterization proposes an intuitive model for CHEs in acid sensing and response via CFTR mediated HCO3^-^ secretion, facilitated by long basal processes that interface with submucosal neurons to relay information about luminal acidity, resulting in acidification of the proximal small intestine in the absence of CFTR in ΔF508 CF disease. We provide compelling evidence that CHEs play an important role in CF intestinal pathogenesis.

## Acknowledgements

The authors thank Dr. Michael Goy for providing UGN antibodies and Dr. Peggy Myung (Yale School of Medicine) for technical support with FISH.

## Grants

This project was supported by NIH R01 DK 077065-11 to NA, CFRI grant # 0010230 to NA and the Yale Liver Center award NIH P30 DK034989 Morphology core.

## Disclosures

No conflicts of interest, financial or otherwise are declared by the authors.

## Author Contributions

DCR, JJ, NAA conceived and designed research; DCR, JJ, CM, AS, EZ, ZDS, and KS performed experiments; DCR, JJ, AS, and KS analysed data; MD and DP provided ΔF508 and CF tissue samples; DCR, JJ, AS, KS, and NAA interpreted results of experiments; DCR, JJ, AS, and KS prepared figures; DCR, JJ, and NAA drafted, edited and revised the manuscript.

## Notes

### Competing Interest Statement

The authors have declared no competing interest.

